# Cartilage-penetrating hyaluronic acid hydrogel preserves tissue content and reduces chondrocyte catabolism

**DOI:** 10.1101/2022.05.17.492335

**Authors:** Michael A. Kowalski, Lorenzo M. Fernandes, Kyle E. Hammond, Sameh Labib, Hicham Drissi, Jay M. Patel

## Abstract

Articular cartilage injuries have a limited healing capacity and, due to inflammatory and catabolic activities, often experience progressive degeneration towards osteoarthritis. Current repair techniques generally provide short-term symptomatic relief; however, the regeneration of hyaline cartilage remains elusive, leaving both the repair tissue and surrounding healthy tissue susceptible to long-term wear. Therefore, methods to preserve cartilage following injury, especially from matrix loss and catabolism, are needed to delay, or even prevent, the deteriorative process. The goal of this study was to develop and evaluate a cartiage-penetrating hyaluronic-acid (HA) hydrogel to improve damaged cartilage biomechanics and prevent tissue degeneration. At time zero, the HA-based hydrogel provided a 46.5% increase in compressive modulus and a decrease in permeability after simulated degeneration of explants (collagenase application). Next, in a degenerative culture model (interleukin-1 β [IL-1β] for 2 weeks), hydrogel application prior to or midway through the culture mitigated detrimental changes to compressive modulus and permeability observed in non-treated explants. Furthermore, localized loss of proteoglycan was observed in degenerative culture conditions alone (non-treated), but hydrogel administration significantly improved the retention of matrix elements. Finally, NITEGE staining and gene expression analysis showed the ability of the HA gel to decrease chondrocyte catabolic activity. These results highlight the importance of reinforcing damaged cartilage with a biomaterial system to both preserve tissue content and reduce catabolism associated with injury and inflammation.

## 1. Introduction

Articular cartilage injuries primarily occur through either traumatic or degenerative events, often creating focal defects in the cartilage surface [1]. The relative avascularity of cartilage leaves it incapable of self-repair, often necessitating intervention following injury. Post-injury, increased stresses and inflammation often lead to upregulation of catabolic genes in chondrocytes and increased enzymatic degradation from matrix proteases [2–4]. These events have a pronounced impact on reducing the inherently robust mechanical integrity of cartilage [5], which in healthy tissue is a result of a dense extracellular matrix rich in proteoglycans [6]. The high fluid pressurization potential of healthy cartilage in compromised following injury and matrix degradation [7], leaving the tissue susceptible to wear and further cell-mediated catabolism. The resulting degenerative cycle often develops into whole-joint osteoarthritis, which affects over 300 million people worldwide [8]. Thus, there is a clear need to develop novel techniques and methods to prevent, slow, or even reverse this degenerative cascade.

Currently, cartilage injuries are surgically treated through four primary means; removal of the damaged tissue, repair via direct fixation, marrow stimulation, and lastly via transplantation utilizing autograft or allograft tissue [9]. However, these treatments are relatively ineffective in long-term outcomes and/or provide only short term symptomatic relief [10]. Also, the repair tissue often lacks the mechanical rigidity of healthy hyaline cartilage, leaving it vulnerable to long-term wear [9,11,12]. Due to the challenges in hyaline cartilage regeneration, slowing or even preventing progressive deterioration of cartilage is of significant value to the field. Several groups have utilized biomaterials and crosslinking methods to “stabilize” existing cartilage, focusing on biomechanical fortification of the tissue. Specifically, genipin crosslinking can improve the mechanical properties of degraded cartilage, but at the expense of chondrocyte viability [13]. Other techniques have shown that both natural and synthetic hydrogels could be used to resurface [14] or interpenetrate [15–17] damaged cartilage, improving the biphasic mechanical properties of degenerated cartilage tissue at time-zero. However, to our knowledge, the response of chondrocytes past this initial application, especially in inflammatory conditions that are often present post-injury, is relatively unexplored. Thus, there is a need to establish cartilage-fortifying strategies that not only provide initial reinforcement, but also provide prolonged preservation of the biomechanical and biochemical health of cartilage.

Previously, a similar hyaluronic acid (HA) hydrogel biomaterial diffused into a defected cartilage, improved initial mechanics of degenerated cartilage, and remained for at least 7 days in vivo [17]. In the present study, we evaluated a HA-based hydrogel application to reinforce surface damaged cartilage and prevent catabolic deterioration. Specifically, this study confirmed hydrogel interpenetration with cartilage tissue and improved biphasic mechanical properties of degenerated tissue after application. In a degenerative culture model, HA application mitigated biomechanical and biochemical loss and reduced chondrocyte catabolic response. These results present evidence of both time-zero and prolonged benefits from an HA-based cartilage stabilization strategy to potentially delay the progression of cartilage deterioration.

## 2. Materials and Methods

### 2.1 Hyaluronic Acid Synthesis

Methacrylated hyaluronic acid (MeHA; **Fig 1A**) was synthesized from sodium hyaluronate (75kDa; LifeCore Biomedical) and methacrylic anhydride [18,19]. Sodium hyaluronate was dissolved in deionized water (10 mg/mL, 1% w/v), to which a 20-fold excess of methacrylic anhydride was added. The reaction was maintained at a pH of 8.0-9.0 for 6 hours at 4°C. At the end of the reaction, the solution was stirred vigorously overnight at room temperature to degrade the methacrylic anhydride. The solution was dialyzed (MWCO: 6,500) for 10 days, followed by freezing at -20°C for 6 hours and lyophilization (−50°C, 0.05mbar) for 5 days to produce dry MeHA (100% modification confirmed with nuclear magnetic resonance [NMR]; Supplementary Figure 1). Throughout this study, MeHA solution (4% w/v, unless otherwise noted) with LAP photo-initiator (0.05% w/v) was applied to the surface of cartilage explants, given 5-10 minutes to diffuse and photo-crosslinked (3 minutes, 25mW/cm^2^; **Fig 1B**).

**Fig 1.**
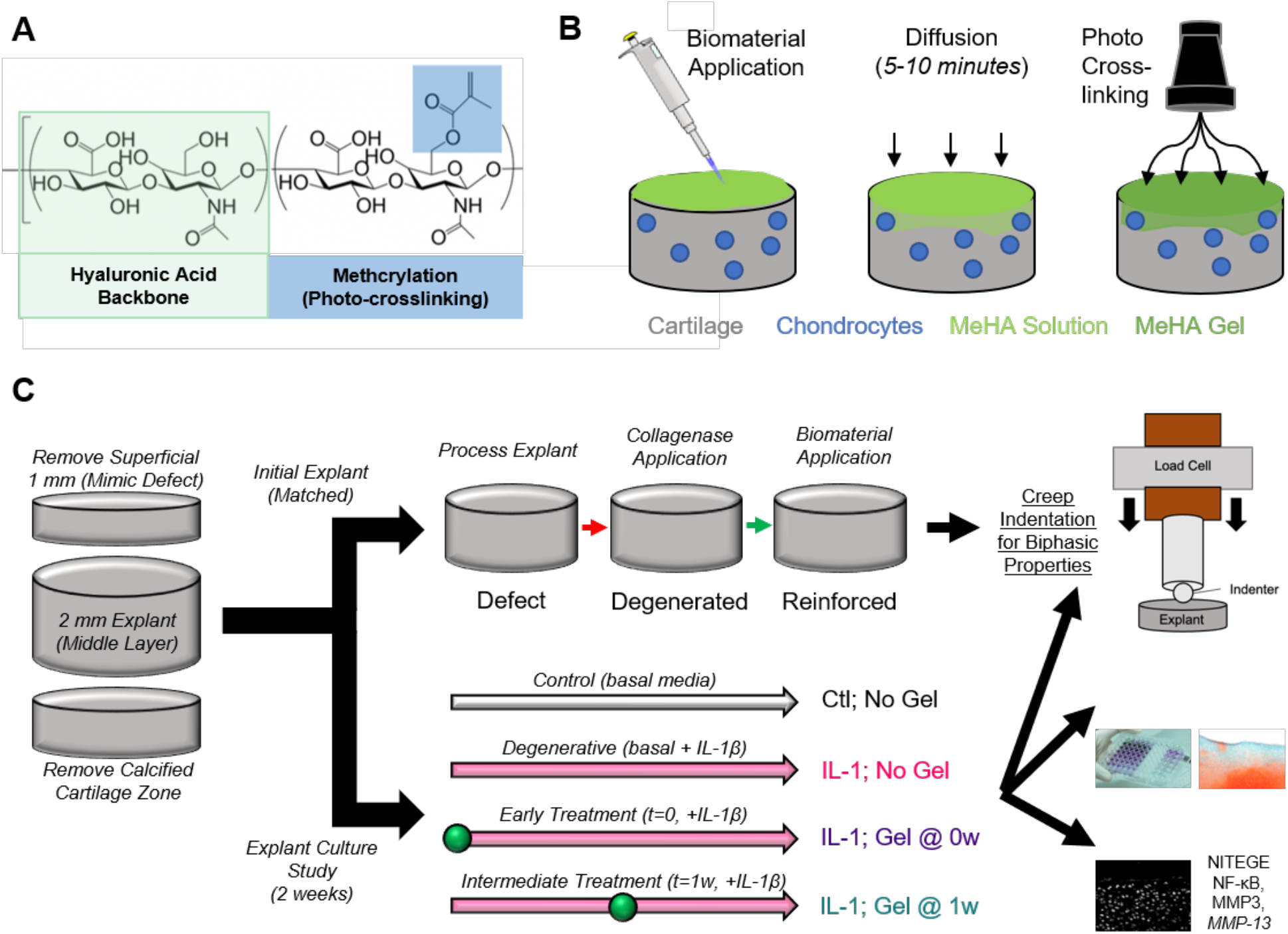
Biomaterial Design, Application, and Study Overview. A) Methacrylated hyaluronic acid backbone to enable photocrosslinking via UV light. B) Schematic depicting biomaterial solution applied to surface of explant, diffusion into cartilage tissue, and photocrosslinking to form interpenetrating gel. C) Study design depicting processing of explants and outline for the initial explant (time-zero mechanics) and degenerative culture studies.

### 2.2 Explant Dissection and Processing

Juvenile bovine (3-6 weeks; Research 87; Boylston MA) cartilage explants (full thickness, 6mm diameter) were harvested from the trochlear groove. After harvest, the calcified cartilage region was removed, leaving the top 3mm of cartilage tissue. Next, the superficial 1mm of cartilage was removed to create a consistent cartilage defect model (**Fig 1C**). To verify diffusion and retention of the hydrogel in the explants, methacrylated rhodamine (0.025% w/v) was added to the MeHA solution, allowing for visualization of hydrogel interdigitation. After verification of gel interpenetration, functional outcome metrics in two studies (initial explant and degenerative culture) were conducted to assess the mechanical properties, cartilage health, and the efficacy of reinforcement of the cartilage tissue (**Fig 1C**).

### 2.3 Time-Zero Explant Mechanical Testing

We performed an initial study to assess the reinforcement that the MeHA hydrogel provides to degenerated cartilage explants. Cartilage explants were subjected to sequential Hertzian Indentation creep tests following processing (defect condition), degeneration (digestion in 0.1% collagenase for 45 minutes), and biomaterial application (digested explant reinforced with MeHA). Creep testing involved applying a 0.25N constant load (Biomomentum Mach-1) on the explant surface for 15 minutes with a spherical indenter (r=2mm), which was fit with a Hertzian biphasic creep model [20] to determine the compressive modulus and permeability (**Fig 1C**).

### 2.4 Degenerative Culture

To emphasize media exposure to the surface of explants, we utilized agarose wells to confine cartilage samples. We added 1 mL of sterile, liquid agarose to each well of a 24-well non-treated tissue culture plate, and allowed cooling until gel formation. A 6mm biopsy punch was then used to excise agarose to create a well matched to the size of explants. Explants were then treated with either basal media as a control (Dulbecco’s Modified Eagle’s Medium [DMEM], 10% Fetal Bovine Serum [FBS], and 1% penicillin-streptomycin-fungizone [PSF]) or basal media supplemented with 10 ng/mL of IL-1β (human recombinant; Tonbo Bioscience 218018U010) for 2 weeks. In addition to a degenerated condition alone (IL-1β with no MeHA), and two MeHA application windows were tested. To mimic a focal defect scenario (with little surface degeneration), MeHA was applied at the start of the study (t0), followed by 2 weeks of culture in IL-1β). To mimic a more intermediate state of degeneration, MeHA was applied one-week into IL-1β culture (t1), and cultured for an additional week in IL-1β. After the conclusion of the culture, explants were subjected to the Hertzian creep indentation testing described previously, as well as glycosaminoglycan (GAG) quantification, histological analysis, immunofluorescence staining, and gene expression (**Fig 1C**).

### 2.5 DMMB Assay

To determine sulfated GAG (s-GAG) content of cultured explants, a dimethylmethylene blue (DMMB) assay was performed. At 2 weeks of culture, explants were digested in buffered papain solution (0.1M Sodium Acetate (anhydrous), 10mM Cysteine Hydrochloride, 50mM EDTA, pH 6, 2% v/v papain) at 25mg dry tissue per 1mL papain solution (48 hours at 60°C). Following digestion, the solution was plated in duplicate in a 96-well assay plate (5uL), followed by addition of 200uL of 1,9-dimethylmethylene blue (8mg DMMB, 2.5mL ethanol, 1g sodium formate, 1mL formic acid, and 496.5mL deionized water). Absorbance values at 525nm were quantified relative to a standard curve of chondroitin sulfate (Sigma C-4384). s-GAG content was normalized as mass per dry weight of tissue.

### 2.6 Safranin-O Fast Green Staining

To determine spatial proteoglycan content, especially relative to the tissue surface, Safranin-O Fast Green staining was performed. Solutions of 0.05% w/v Fast green, 0.1% w/v Safranin O, and 1% v/v acetic acid were used. After mechanical testing, cartilage explants were fixed in formalin, frozen in optical cutting temperature (OCT) compound, sectioned at 20µm, and rinsed twice with DI H_2_O. The sections were then sequentially submerged for 5 minutes in Fast Green (0.05% w/v), 20 seconds in acetic acid (1% v/v), and 10 minutes in Safranin O (0.1% w/v), followed by 2 tap water rinses, dehydration (95%, 100%, 2X each), clearing with Xylene, and mounting and coverslipping.

To quantify the Safranin-O images, a custom written MATLAB code was developed. The code first inputs the grayscale image of red pixels from the original histology image. Next, the code allows the user to create 5 lines from the surface of the explant through the depth of the tissue. The code then takes average pixel intensities (the 10 nearest pixels) every micron along each of the 5 lines, until a depth 500µm. Thus, the code plots the average red pixel intensity (indicative of remaining proteoglycan) as a function of depth from the cartilage surface.

### 2.7 Immunofluorescence Staining

Additional 20µm histological sections were rinsed (3X PBS). Antigen retrieval was performed via digestion with Proteinase K (0.1% v/v) for 5 minutes, followed by 3X PBS rinses. Next, to prevent non-specific binding, a blocking step with 1% bovine serum albumin (BSA) in PBS was applied for 30 minutes. Primary antibodies were applied in 1% BSA for 1 hour at room temperature, and included type VI collagen(rabbit monoclonal; Abcam ab182744; 1:200) for detection of the pericellular matrix during interdigitation verification and NITEGE (rabbit polyclonal; Fisher PA1-1746; 1:200) for detection of aggrecan breakdown via aggrecanases [21,22]. Following 3x PBS rinses, sections were stained with secondary antibody in 1% BSA for 1 hour, rinsed 3X PBS, and mounted with Prolong Gold Antifade Reagent with DAPI and cover-slipped. To quantify the relative NITEGE staining between groups, a histogram plot of the intensity values was plotted in FIJI, quantifying the intensity and area of NITEGE positive staining [23].

### 2.8 Gene Expression

Another set of cartilage explants were harvested for gene expression of catabolic markers. The top ∼500µm of tissue was placed in TRIzol (Invitrogen) for RNA isolation according to the manufacturer’s protocol (the remaining tissue was discarded). Following RNA quantification, cDNA was synthesized using oligo(dT) and random primers with qScript cDNA SuperMix (Quantabio). Finally, the Applied Biosystems 7500 Real-Time PCR System was used with PowerUP SYBR green master mix (Applied Biosystems) to perform all qPCR reactions. Melt curve analysis was performed to confirm amplicon authenticity and the ΔΔCt method [24] was used for fold change analyses. Expression of MMP-13, MMP-3, and NF-κB were analyzed, with β-actin serving as a housing keeping gene.

### 2.9 Statistical Analysis and Rigor

Explants from at least 3 donors were included for each analysis to account for donor variability. All data was subject to outlier (ROUT method) and normality (Shapiro-Wilk) testing. Parametric, normal datasets were analyzed with a one-way analysis of variance (ANOVA) with post-hoc Tukey’s testing. Nonparametric or non-normal datasets were analyzed with a Kruskal-Wallis test with post-hoc Dunn’s Multiple Comparison Test. All data are shown as dot plots for transparency, and p<0.05 was chosen as a threshold for statistical significance.

## 3. Results

### 3.1 Time-Zero Interpenetration and Reinforcement

Fluorescence imaging showed retention of the gel in the first few hundred µm of explants (**Fig 2A**). Upon closer examination, the MeHA gel showed a greater localization to the area around each chondrocyte, as depicted by type VI collagen, a marker of the chondrocyte pericellular matrix (PCM) (**Fig 2B**). The MeHA gel diffuses an average of 156.39µm from the surface of application (Supplementary Figure 2). Next, defected cartilage explants (“Defect”) were tested with indentation creep testing, showing some variability in compressive modulus (112 – 376 kPa). For this reason, sequential testing after both degeneration and reinforcement was performed. Degeneration in collagenase led to a 19.5% decrease in compressive modulus (p = 0.0078); reinforcement with the MeHA gel following this simulated degeneration led to a 46.5% increase over their prior degenerated state (p = 0.082) (**Fig 2C**). Permeability values showed an opposite trend, with an increase after degeneration and recuperation after reinforcement (Defect vs. Degenerated - p = 0.0029; Degenerated vs. Reinforced - p = 0.0065) (**Fig 2D**). As a result of improved mechanical properties following gel application, other concentrations of the MeHA gel (1, 2% w/v) were also tested. Both 1% and 2% gel application led to partial recovery of compressive modulus, while only the 2% application led to partial recovery of permeability (Supplementary Figure 3). For comparison, we took the average loss in compressive modulus following degeneration and determined how much of this loss was recovered by reinforcement. The 4% gel showed the highest recovery in compressive modulus (**Fig 2E**); thus, the 4% gel was used for the degenerative culture model.

**Fig 2.**
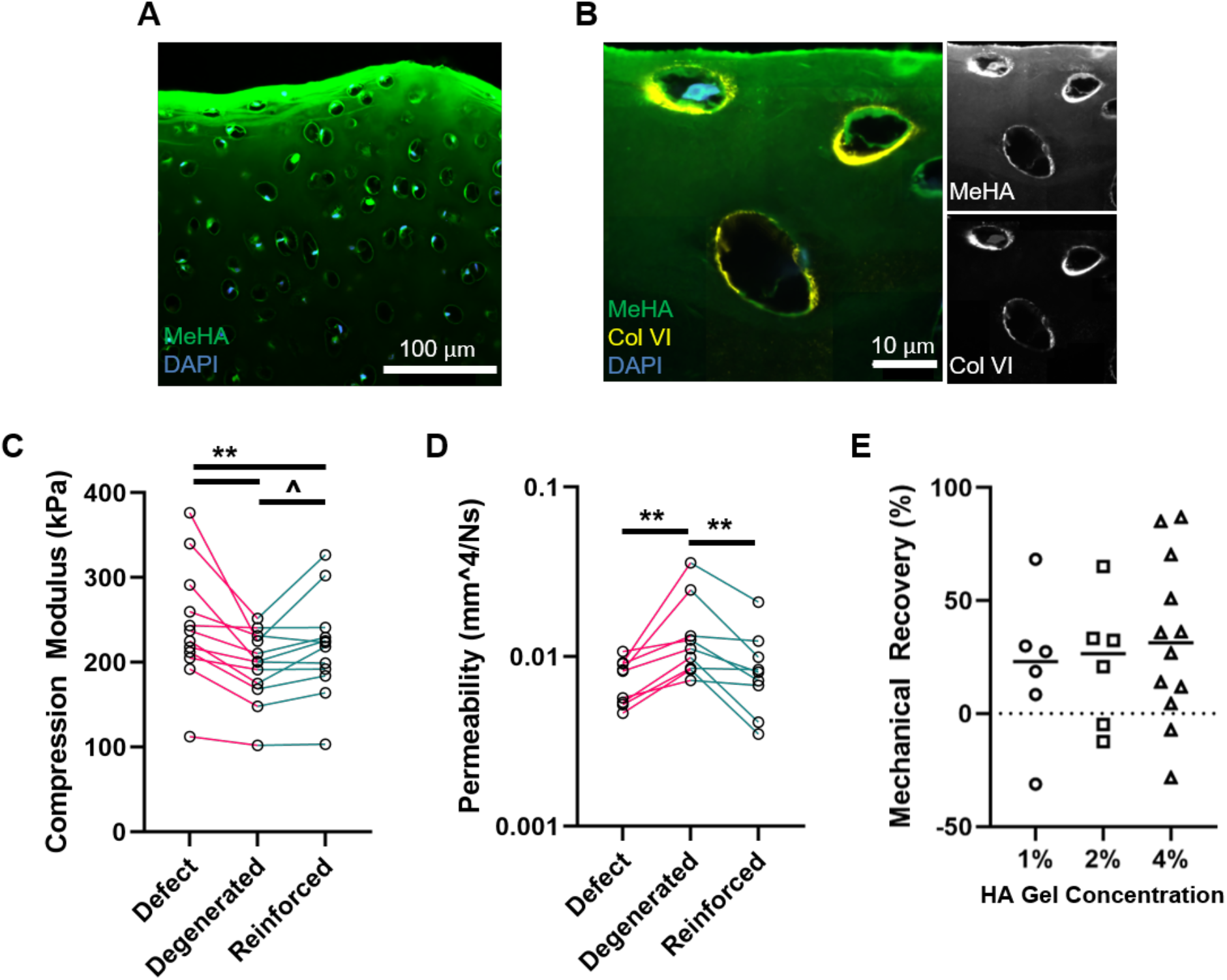
Biomaterial Interdigitation and Mechanical Reinforcement. A) Immunofluorescent image of MeHA gel (green) interpenetrating with cartilage tissue (chondrocyte nuclei in blue; DAPI). Scale bar = 100µm. B) Immunofluorescent image of MeHA gel (green) colocalizing with chondrocyte pericellular matrix shown via a type VI collagen stain (yellow). Scale bar = 10µm. Insets show MeHA and type VI collagen individually. C) Compressive modulus and D) permeability of cartilage tissue in defected, degenerated, and reinforced scenarios. n=12 per group. E) Mechanical recovery of compressive modulus, defined by the percentage modulus recovered after reinforcement (1, 2, 4% MeHA; n=6,6,12 per group, respectively) of degenerated explants. ^, *, ** represent p<0.10, 0.05, and 0.01, respectively.

### 3.2 Tissue Preservation in Degenerative Culture

Next, we investigated the cartilage-preserving aspects of the MeHA biomaterial in a two-week inflammatory culture. Cell viability of post processed explants and application of biomaterial was assessed following 24 hours of basal media culture to ensure significantly different viability was not observed following biomaterial application (Supplementary Figure 4). A significant decrease (67.8%, p<0.0001) in compressive modulus was observed as a result of IL-1β treatment; however, MeHA gel application at the start of inflammatory culture mitigated (52.5% recovery, IL-1 vs t0; p = 0.0125) much of this mechanical loss (**Fig 3A**). MeHA application at 1-week into the inflammatory culture produced a trend (36.4% recovery, IL-1 vs t1; p = 0.1277) in retainment of compressive modulus. Similar trends were seen with regards to permeability (**Fig 3B**), with worsening permeability (increased permeability; p = 0.0002) with IL-1β treatment alone, and mitigation of this increase with both t0 (p = 0.0128) and t1 (p = 0.023) MeHA application.

**Fig 3.**
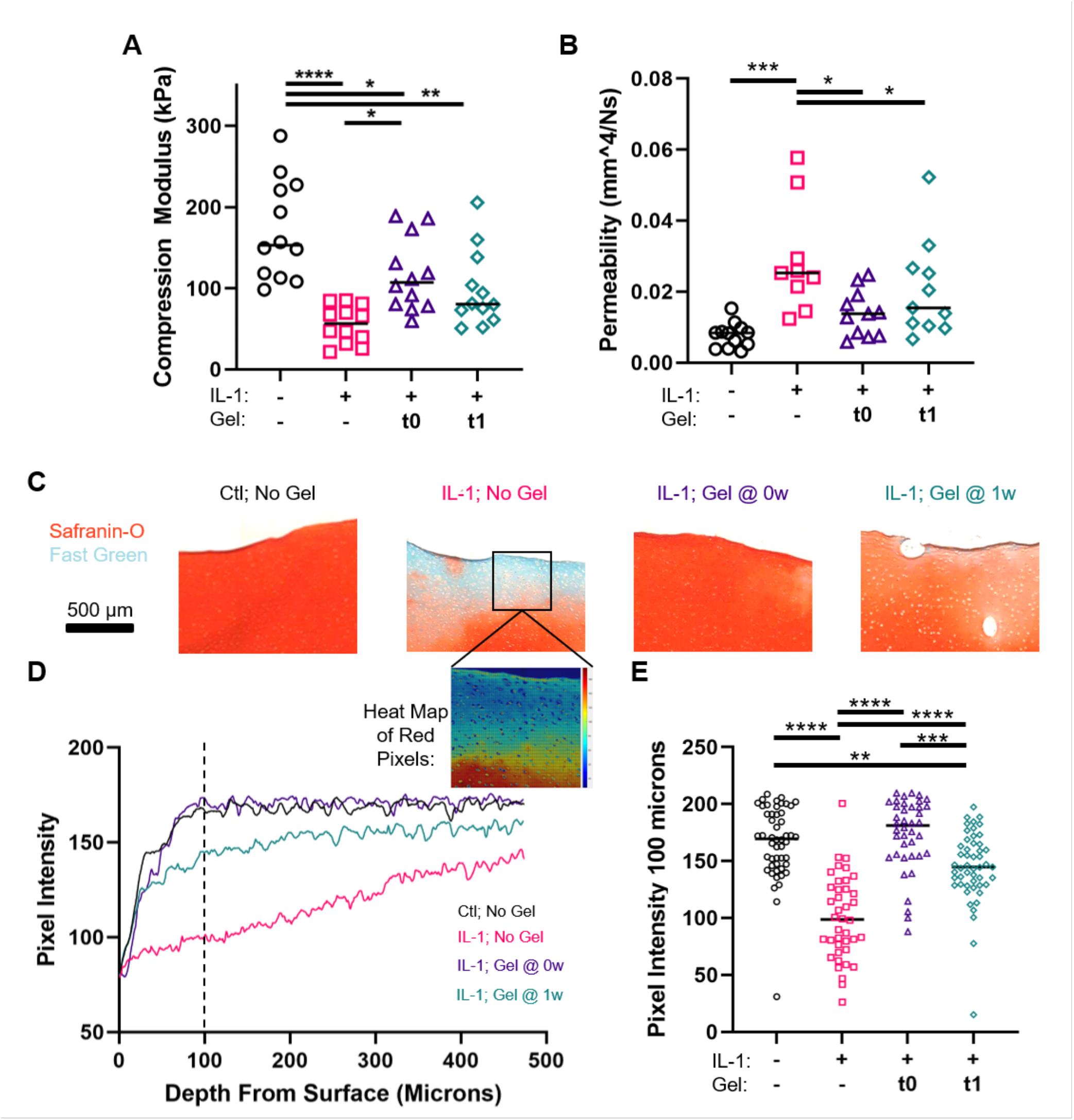
Cartilage Tissue Preservation in Degenerative Culture via MeHA Gel. A) Compressive modulus and B) permeability of explants cultured in control media, IL-1β media (no gel), and IL-1β media with the 2 MeHA gel application points (t0, t1). n=12 per group. C) Safranin O-Fast Green stained sections from the 4 groups. Scale Bar = 500µm. D) Quantification of red channel intensity as a function of depth from the surface (average profile shown, n=8 per group). E) Bar graph of red pixel intensity 100µm from the cartilage surface (n=8 per group). *, **, ***, **** represent p<0.05, 0.01, 0.001, and 0.0001, respectively.

To assess spatial and bulk biochemical changes in explants, Safranin O-Fast Green staining and a DDMB assay were performed. In comparison to healthy proteoglycan content in controls indicated by a deep red stain, after inflammatory culture explants showed a loss of Safranin O staining, particularly at the surface exposed to media (**Fig 3C**). Application of the MeHA biomaterial (t0 and t1) led to the retention of red proteoglycan staining, with a slight observable loss of red staining in the t1 explants. Biomaterial application at time-zero showed the potential for reduced biochemical degradation in an extreme inflammatory state (20 ng/mL IL-1β for 3 weeks, Supplementary Figure 5). Quantification of the pixel intensity as a function of depth confirmed these findings, where application of the biomaterial at t0 and t1 timepoints led to complete and partial retention, respectively, of proteoglycan content in the first ∼300µm of the tissue (**Fig 3D**). Five 500µm lines were selected for each image and each pixel on the line was averaged with the ten pixels around it to achieve a pixel intensity metric by depth (Supplementary Figure 6). Red pixel intensity at 100µm from the surface showed a significant decrease from control with IL-1 treatment (p-value < 0.0001), but a retention of pixel intensity following application of explants at both time points compared to IL-1 treatment (p-values < 0.0001). Control explants and application prior to inflammatory culture showed no significant difference in pixel intensity indicating proteoglycan retention similar to control explants (p-value > 0.9999). Application timepoints prior to culture and one week in to culture showed a significant difference in proteoglycan positive staining (p-value = 0.0008). Also, application one week into inflammatory culture showed a significant red pixel intensity decrease from culture explants (p-value = 0.0076). The s-GAG content in entire explants showed no significant differences between groups (Supplementary Figure 7).

### 3.3. Matrix and Chondrocyte Catabolism

To better characterize signs of matrix and cell catabolism, we investigated key markers of aggrecan degradation (NITEGE) and multiple catabolic markers *NF-κB, MMP-3* and MMP-13 downstream of IL-1β [21,25]. Exposure to IL-1β for 2-weeks showed a drastic increase in NITEGE staining, which was nearly completely diminished by MeHA gel application at either timepoint (**Fig 4A**). Quantification of NITEGE showed a significant reduction of the number of high-intensity pixels (Intensity: 200-255) in gel application samples compared to IL-1β stimulated samples (**Fig 4B,C**). To further explore the catabolic activity occurring at the cellular level, gene expression analysis of *MMP-13, NF-κB*, and *MMP-3* were conducted. IL-1β exposure significantly increased total *MMP-13* and *NF-κB* expression (26.86-and 4.2-fold, p< 0.0001 and p = 0.0005, respectively) (**Fig 4D, F**). MeHA gel application provided a significant reduction in expression of *NF-κB* (0.363-and 0.347-fold, for both conditions t0 and t1, p = 0.0023) (**Fig 4D**) and a trend of reduction in expression of *MMP-13* (0.457-and 0.448-fold, p = 0.0976 and p = 0.1950, for t0 and t1, respectively) (**Fig 4F**). *MMP-3* showed a significant increase (p=0.0035) in expression in chondrocytes when stimulated with IL-1β and MeHA application reduced *MMP-3* expression, albeit not significantly (t1 application, p = 0.1103) (**Fig 4E**).

**Fig 4.**
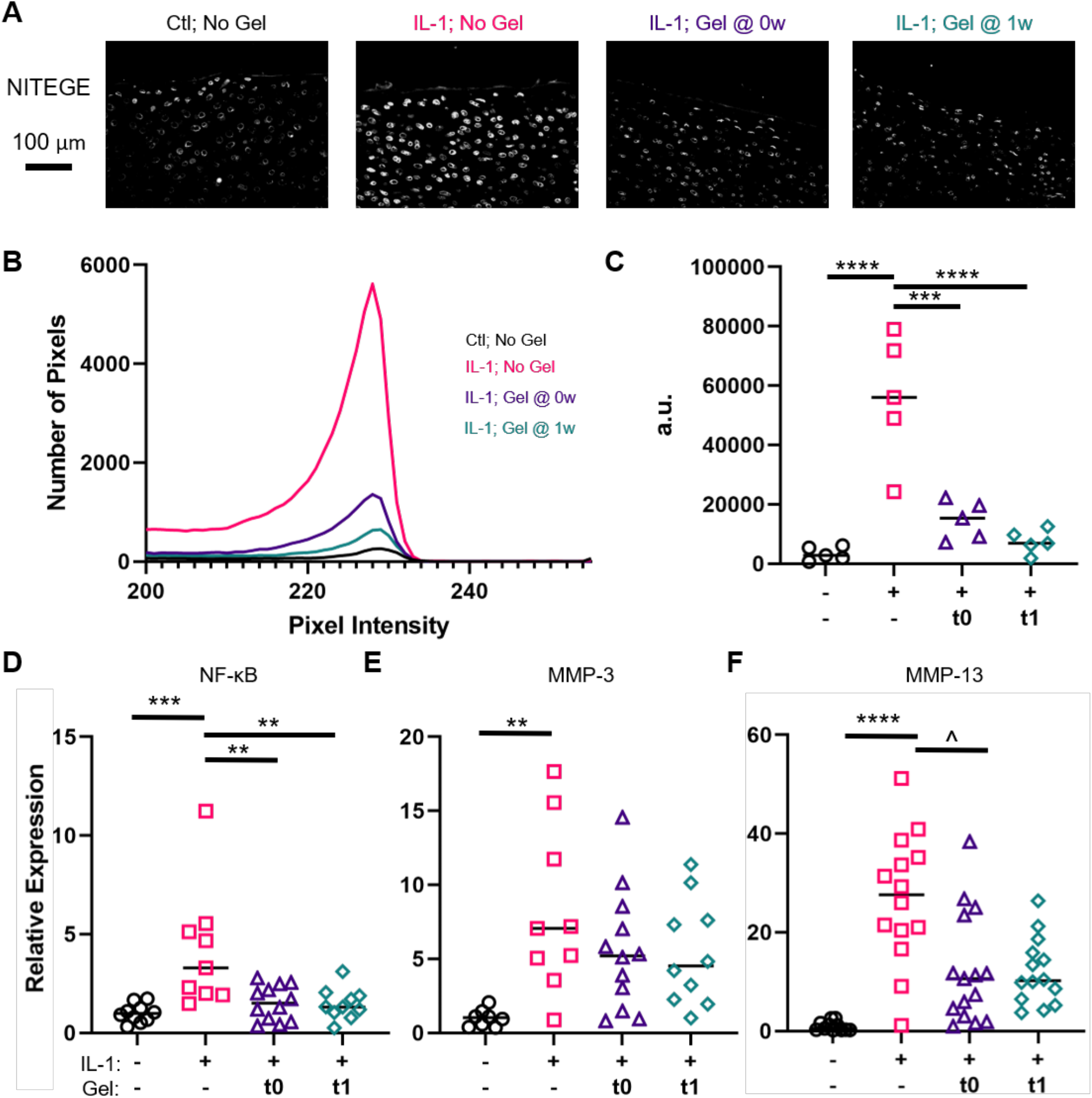
Catabolic Attenuation in Degenerative Culture. A) Immunofluorescence staining of NITEGE neo-epitope (aggrecan degradation) in the 4 groups. Scale bar = 100µm. B) Quantification of NITEGE pixel intensity (average profile shown, n=5 per group) from grayscale value 200-255. C) Quantification of the area under each line graph of NITEGE quantification in arbitrary units (pixels*intensity; n=5 per group). D) NF-κB, E) MMP-3, and MMP-13 gene expression (top 500µm of explants) in the 4 groups. n=14 per group. ^, **, ***, **** represent p< 0.1, 0.01, 0.001, 0.0001, respectively.

## 5. Discussion

Due to the progressive deterioration that cartilage injuries often undergo, it is imperative to identify solutions that stabilize the damaged tissue and prevent degeneration that results from inflammation, catabolic activities, and compromised joint loading [26]. Fortifying the injured tissue as early as possible could provide increased stability and restore mechanical properties (including fluid pressurization via low permeability) to near-healthy levels. This would also translate to less stress on chondrocytes in the injured tissue, potentially preventing the production of catabolic factors in stress-elevated environments [27,28]. While time-zero mechanical properties of cartilage tissue have been improved with various techniques, the response of cells within and maintenance of tissue properties have not, to our knowledge, been reported. In the present study, we verified the ability of a MeHA hydrogel to interdigitate with and reinforce degenerated cartilage [17]. Further, in a degenerative cartilage explant model, we demonstrated the maintenance of matrix elements and reduced catabolic cell activity [29]. Overall, this initial reinforcement, combined with lower tissue degradation and chondrocyte catabolism, could offer a method to delay, or even prevent, the post-injury deteriorative process.

Penetration on small molecules into cartilage tissue is of great interest in treatment modalities. Previous literature shows nanoparticles can diffuse into cartilage tissue and present an opportunity for using cartilage penetration strategies to deliver bioactive factors [30]. Other diffusing materials have provided a mechanical benefit to the damaged tissue [16,31]. Through fortification strategies, researchers have provided increased stability and restored mechanical properties (including fluid pressurization via low permeability) to near-healthy levels. Polyethylene glycol (PEG) hydrogels of varying concentrations have been shown to improve the mechanical modulus of cartilage from 30-286% [16]. Genipin crosslinking has also been shown to improve the mechanical modulus of cartilage anywhere from 20-80% post treatment [32]. The present study similarly restored, at least partially, the compressive modulus of cartilage that is compromised after degeneration, also shown previously with MeHA hydrogels [17]. Recovery of low permeability, similar to native cartilage, was also observed after hydrogel application.

This study then investigated the continuing effects of hydrogel reinforcement on cartilage biochemistry and biomechanics in a severe inflammatory state, a new line of experiments not performed by prior cartilage reinforcement techniques. Increased inflammation in the synovium and joint often accompany cartilage and other joint injuries [33]. As a result, these inflammatory factors (e.g. IL-1β) often contribute to further degradation of the tissue and its function [34]. Previous *in vitro* degeneration models have shown similar loss of proteoglycan content and Young’s modulus, and an upregulation of catabolic activities through IL-1β treatment [35–37]. In the degenerative culture environment in this study, the interpenetrating HA hydrogel preserved the mechanical integrity and matrix content of the tissue. This could be the result of stabilization of the tissue in the initial injury state and sustained reinforcement of the tissue to prevent extreme degradation resulting from injury, inflammation, or increased loading stress. This reinforcement may also provide protection of cell activity from catabolism through specific reinforcement of the PCM around chondrocytes [29,38], positively benefiting their mechano-transduction.

Another potential contributing factor to both biomechanical and biochemical benefit in the degenerative culture is hyaluronic acid signaling, which has been shown to have an anti-inflammatory effect in cartilage tissue [39]. In healthy tissue, HA is an extremely abundant glycosaminoglycan and provides lubrication in the joint synovium via lubricin, making it vital to cartilage health [40]. Furthermore, chondrocytes interact with HA through CD44 signaling, which is proven to affect cell function and chondrogenesis [41,42]. Injections of hyaluronic acid have proven inconsistent in providing a therapeutic effect for patients with osteoarthritis; this is likely attributed to the fast clearance out of the joint [43]. Our study provides an option for sustained HA therapy through crosslinking of a hydrogel within damaged tissue, potentially provide anti-inflammatory signaling to the chondrocytes exposed to a degenerative environment. The biomechanical benefits observed by applying gel can likely be attributed to a biochemical benefit in terms of localized s-GAG retention in the surface of explants (where both degenerative media and hydrogel application are focused). This anti-inflammatory effect was also observed at the cellular level with NITEGE staining (neoepitopes of aggrecan degradation), which showed significant staining in the inflammatory condition, but was drastically reduced in application conditions. Gene expression of surface chondrocytes confirmed this trend, with NF-κB (downstream of IL-1β signaling) expression significantly reduced and MMP-13 showing a trend of reduction in hydrogel application groups.

In the present study, hydrogel interdigitation with cartilage provided promise in preventing degradative changes, potentially due to both reinforcement and anti-catabolic effects. In the future, the biomaterial can be uniquely tailored to patients due to the easily modifiable nature of HA. Specifically, the level of fortification can be readily modified by changing the degree of modification, gel concentration, and photocrosslinking parameters [44]. As a result, gels can be formulated to treat various injury types that would require a specific degree of reinforcement. Another aspect worth exploration includes understanding how the MeHA gel application affects cell behavior and interacts with the extracellular matrix (ECM) and PCM. Specifically, localization of the material to the PCM brings new questions regarding whether the observed protective effect is through reinforcement of PCM mechanics, through prolonged HA signaling, or perhaps both. Thus, future work will aim to study the mechanism behind the protective effects of our MeHA hydrogel, as well as translate this approach to animal models for functional evaluation.

The results of this study clearly demonstrate that a cartilage-interdigitating MeHA hydrogel can reinforce the tissue initially and mitigate cartilage degeneration. Certainly, there were a few limitations regarding our approach: *in vitro* nature of all assays, simple degenerative model, and juvenile bovine cartilage explants. *In vitro* studies fail to replicate the actual conditions that occur in the cartilage and joint injuries that could alter the effect of the application. A simple degenerative model using only one inflammatory factor also fails to recapitulate the joint and cartilage lesion environment post injury due to the multitude of inflammatory factors that are present in the joint and synovium. This is certainly seen as a next step, to evaluate the therapy in a more complex degenerative explant model, and even in a small animal cartilage defect study. Finally, variability between donors can lead to a large range of mechanical properties and responses to degenerative culture/ treatment present another limitation when using an explant model, though these concerns were mitigated by performing analyses in at least 3 donors per assay.

## 5. Conclusion

In summary, we demonstrated that MeHA hydrogel provides initial mechanical benefits to damaged cartilage and mitigates tissue loss in an inflammatory culture. We also demonstrated that the MeHA biomaterial provides matrix-protective effects of effectors downstream of IL-1β like MMP-13 and NF-κB, while reducing the expression of aggrecanase products. As a result, we believe that this MeHA hydrogel application system is a promising biomaterial approach to delaying the onset of osteoarthritis.

## Abbreviations

HA: Hyaluronic Acid
IL: Interleukin
NITEGE: C-terminal neoepitope
MeHA: Methacrylated hyaluronic acid
kDa: kilodalton
w/v: weight per volume
MWCO: molecular-weight cutoff
NMR: nuclear magnetic resonance
LAP: Lithium phenyl-2,4,6-trimethylbenzoylphosphinate
mW: milli Watts
PBS: phosphate-buffered saline
UV: ultraviolet
DMEM: Dulbecco’s Modified Eagle’s Medium
FBS: Fetal bovine serum
PSF: penicillin-streptomyscin-fungizone
GAG: glycosaminoglycan
s-GAG: sulfated glycosaminoglycan
DMMB: dimethylmethylene blue
EDTA: Ethylenediaminetetraacetic acid
OCT: optimal cutting temperature
BSA: bovine serum albumin
DAPI: 4′,6-diamidino-2-phenylindole
RNA: ribonucleic acid
DNA: deoxyribonucleic acid
MMP: matrix metalloproteinase
ROUT: regression and outlier
ANOVA: analysis of variance
PCM: pericellular matrix
PEG: polyethylene glycol
ECM: extracellular matrix

## 6. Acknowledgements

The authors would like to acknowledge the Emory Department of Orthopaedics and the Emory NMR Facility for their assistance with this work. The authors also thank Drs. Johnna Temenoff and Andres Garcia for their input. This work was funded by the Department of Veterans Affairs [IK1 RX003208, IK2 RX003928] and startup funds from the Emory University School of Medicine.

## References

[1] J.A. Buckwalter, Articular cartilage injuries, in: Clin. Orthop. Relat. Res., 2002. https://doi.org/10.1097/00003086-200209000-00004.

[2] T.M. Simon, D.W. Jackson, Articular Cartilage: Injury Pathways and Treatment Options, Sports Med. Arthrosc. (2018). https://doi.org/10.1097/JSA.0000000000000182.

[3] D.J. Griffin, J. Vicari, M.R. Buckley, J.L. Silverberg, I. Cohen, L.J. Bonassar, Effects of enzymatic treatments on the depth-dependent viscoelastic shear properties of articular cartilage, J. Orthop. Res. (2014). https://doi.org/10.1002/jor.22713.

[4] Z. Fan, B. Bau, H. Yang, S. Soeder, T. Aigner, Freshly isolated osteoarthritic chondrocytes are catabolically more active than normal chondrocytes, but less responsive to catabolic stimulation with interleukin-1β, Arthritis Rheum. (2005). https://doi.org/10.1002/art.20725.

[5] R.E. Wilusz, S. Zauscher, F. Guilak, Micromechanical mapping of early osteoarthritic changes in the pericellular matrix of human articular cartilage, Osteoarthr. Cartil. (2013). https://doi.org/10.1016/j.joca.2013.08.026.

[6] J.A. Buckwalter, H.J. Mankin, Articular cartilage: tissue design and chondrocyte-matrix interactions., Instr. Course Lect. (1998).

[7] G.A. Ateshian, The role of interstitial fluid pressurization in articular cartilage lubrication, J. Biomech. (2009). https://doi.org/10.1016/j.jbiomech.2009.04.040.

[8] S. Safiri, A.A. Kolahi, E. Smith, C. Hill, D. Bettampadi, M.A. Mansournia, D. Hoy, A. Ashrafi-Asgarabad, M. Sepidarkish, A. Almasi-Hashiani, G. Collins, J. Kaufman, M. Qorbani, M. Moradi-Lakeh, A.D. Woolf, F. Guillemin, L. March, M. Cross, Global, regional and national burden of osteoarthritis 1990-2017: A systematic analysis of the Global Burden of Disease Study 2017, Ann. Rheum. Dis. (2020). https://doi.org/10.1136/annrheumdis-2019-216515.

[9] A.R. Martín, J.M. Patel, H.M. Zlotnick, J.L. Carey, R.L. Mauck, Emerging therapies for cartilage regeneration in currently excluded ‘red knee’ populations, Npj Regen. Med. (2019). https://doi.org/10.1038/s41536-019-0074-7.

[10] A.J. Krych, D.B.F. Saris, M.J. Stuart, B. Hacken, Cartilage Injury in the Knee: Assessment and Treatment Options, J. Am. Acad. Orthop. Surg. (2020). https://doi.org/10.5435/JAAOS-D-20-00266.

[11] A.G. McNickle, M.T. Provencher, B.J. Cole, Overview of existing cartilage repair technology, Sports Med. Arthrosc. (2008). https://doi.org/10.1097/JSA.0b013e31818cdb82.

[12] J.C. Hu, K.A. Athanasiou, The effects of intermittent hydrostatic pressure on self-assembled articular cartilage constructs, Tissue Eng. (2006). https://doi.org/10.1089/ten.2006.12.1337.

[13] C.M. Bonitsky, M.E. McGann, M.J. Selep, T.C. Ovaert, S.B. Trippel, D.R. Wagner, Genipin crosslinking decreases the mechanical wear and biochemical degradation of impacted cartilage in vitro, J. Orthop. Res. (2017). https://doi.org/10.1002/jor.23411.

[14] S. Grenier, P.E. Donnelly, J. Gittens, P.A. Torzilli, Resurfacing damaged articular cartilage to restore compressive properties, J. Biomech. (2015). https://doi.org/10.1016/j.jbiomech.2014.10.023.

[15] J.T.A. Mäkelä, B.G. Cooper, R.K. Korhonen, M.W. Grinstaff, B.D. Snyder, Functional effects of an interpenetrating polymer network on articular cartilage mechanical properties, Osteoarthr. Cartil. (2018). https://doi.org/10.1016/j.joca.2018.01.001.

[16] B.G. Cooper, R.C. Stewart, D. Burstein, B.D. Snyder, M.W. Grinstaff, A Tissue-Penetrating Double Network Restores the Mechanical Properties of Degenerated Articular Cartilage, Angew. Chemie - Int. Ed. (2016). https://doi.org/10.1002/anie.201511767.

[17] J.M. Patel, C. Loebel, K.S. Saleh, B.C. Wise, E.D. Bonnevie, L.M. Miller, J.L. Carey, J.A. Burdick, R.L. Mauck, Stabilization of Damaged Articular Cartilage with Hydrogel-Mediated Reinforcement and Sealing, Adv. Healthc. Mater. (2021) 2100315. https://doi.org/10.1002/adhm.202100315.

[18] K.A. Smeds, M.W. Grinstaff, Photocrosslinkable polysaccharides for in situ hydrogel formation, J. Biomed. Mater. Res. (2001). https://doi.org/10.1002/1097-4636(200101)54:1<115::AID-JBM14>3.0.CO;2-Q.

[19] J.A. Burdick, C. Chung, X. Jia, M.A. Randolph, R. Langer, Controlled degradation and mechanical behavior of photopolymerized hyaluronic acid networks, Biomacromolecules. (2005). https://doi.org/10.1021/bm049508a.

[20] A.C. Moore, J.F. DeLucca, D.M. Elliott, D.L. Burris, Quantifying Cartilage Contact Modulus, Tension Modulus, and Permeability With Hertzian Biphasic Creep, J. Tribol. 138 (2016) 414051–414057. https://doi.org/10.1115/1.4032917.

[21] P.L.E.M. Van Lent, L.C. Grevers, A.B. Blom, O.J. Arntz, F.A.J. Van De Loo, P. Van Der Kraan, S. Abdollahi-Roodsaz, G. Srikrishna, H. Freeze, A. Sloetjes, W. Nacken, T. Vogl, J. Roth, W.B. Van Den Berg, Stimulation of chondrocyte-mediated cartilage destruction by S100A8 in experimental murine arthritis, Arthritis Rheum. (2008). https://doi.org/10.1002/art.24074.

[22] N.A. Zelenski, H.A. Leddy, J. Sanchez-Adams, J. Zhang, P. Bonaldo, W. Liedtke, F. Guilak, Type VI collagen regulates pericellular matrix properties, chondrocyte swelling, and mechanotransduction in mouse articular cartilage, Arthritis Rheumatol. (2015). https://doi.org/10.1002/art.39034.

[23] J. Schindelin, I. Arganda-Carreras, E. Frise, V. Kaynig, M. Longair, T. Pietzsch, S. Preibisch, C. Rueden, S. Saalfeld, B. Schmid, J.Y. Tinevez, D.J. White, V. Hartenstein, K. Eliceiri, P. Tomancak, A. Cardona, Fiji: An open-source platform for biological-image analysis, Nat. Methods. (2012). https://doi.org/10.1038/nmeth.2019.

[24] K.J. Livak, T.D. Schmittgen, Analysis of relative gene expression data using real-time quantitative PCR and the 2-ΔΔCT method, Methods. (2001). https://doi.org/10.1006/meth.2001.1262.

[25] P. Wojdasiewicz, Ł.A. Poniatowski, D. Szukiewicz, The role of inflammatory and anti-inflammatory cytokines in the pathogenesis of osteoarthritis, Mediators Inflamm. (2014). https://doi.org/10.1155/2014/561459.

[26] D.T. Felson, Risk factors for osteoarthritis: Understanding joint vulnerability, in: Clin. Orthop. Relat. Res., 2004. https://doi.org/10.1097/01.blo.0000144971.12731.a2.

[27] S.M. Dai, Z.Z. Shan, H. Nakamura, K. Masuko-Hongo, T. Kato, K. Nishioka, K. Yudoh, Catabolic stress induces features of chondrocyte senescence through overexpression of caveolin 1: Possible involvement of caveolin 1-induced down-regulation of articular chondrocytes in the pathogenesis of osteoarthritis, Arthritis Rheum. (2006). https://doi.org/10.1002/art.21639.

[28] T. Hodgkinson, D.C. Kelly, C.M. Curtin, F.J. O’Brien, Mechanosignalling in cartilage: an emerging target for the treatment of osteoarthritis, Nat. Rev. Rheumatol. 18 (2022) 67–84. https://doi.org/10.1038/s41584-021-00724-w.

[29] A. Karim, A.C. Hall, Chondrocyte Morphology in Stiff and Soft Agarose Gels and the Influence of Fetal Calf Serum, J. Cell. Physiol. (2017). https://doi.org/10.1002/jcp.25507.

[30] A.G. Bajpayee, M. Scheu, A.J. Grodzinsky, R.M. Porter, A rabbit model demonstrates the influence of cartilage thickness on intra-articular drug delivery and retention within cartilage, J. Orthop. Res. (2015). https://doi.org/10.1002/jor.22841.

[31] J.M. Patel, C. Loebel, K.S. Saleh, B.C. Wise, E.D. Bonnevie, L.M. Miller, J.L. Carey, J.A. Burdick, R.L. Mauck, Stabilization of Damaged Articular Cartilage with Hydrogel-Mediated Reinforcement and Sealing, Adv. Healthc. Mater. (2021). https://doi.org/10.1002/adhm.202100315.

[32] M.E. McGann, C.M. Bonitsky, M.L. Jackson, T.C. Ovaert, S.B. Trippel, D.R. Wagner, Genipin crosslinking of cartilage enhances resistance to biochemical degradation and mechanical wear, J. Orthop. Res. (2015). https://doi.org/10.1002/jor.22939.

[33] P. Bhattaram, U. Chandrasekharan, The joint synovium: A critical determinant of articular cartilage fate in inflammatory joint diseases, Semin. Cell Dev. Biol. (2017). https://doi.org/10.1016/j.semcdb.2016.05.009.

[34] J.P. Pelletier, J.A. DiBattista, P. Roughley, R. McCollum, J. Martel-Pelletier, Cytokines and inflammation in cartilage degradation, Rheum. Dis. Clin. North Am. (1993). https://doi.org/10.1016/s0889-857x(21)00331-8.

[35] M. Lv, Y. Zhou, S.W. Polson, L.Q. Wan, M. Wang, L. Han, L. Wang, X.L. Lu, Identification of Chondrocyte Genes and Signaling Pathways in Response to Acute Joint Inflammation, Sci. Rep. (2019). https://doi.org/10.1038/s41598-018-36500-2.

[36] L. Wang, T. Ma, Y. Zheng, S. Lv, Y. Li, S. Liu, Diosgenin inhibits IL-1β-induced expression of inflammatory mediators in human osteoarthritis chondrocytes, Int. J. Clin. Exp. Pathol. (2015).

[37] L. Sun, X. Wang, D.L. Kaplan, A 3D cartilage - Inflammatory cell culture system for the modeling of human osteoarthritis, Biomaterials. (2011). https://doi.org/10.1016/j.biomaterials.2011.04.028.

[38] I.C. Young, S.T. Chuang, A. Gefen, W.T. Kuo, C.T. Yang, C.H. Hsu, F.H. Lin, A novel compressive stress-based osteoarthritis-like chondrocyte system, Exp. Biol. Med. (2017). https://doi.org/10.1177/1535370217699534.

[39] Y. Jin, R.H. Koh, S.H. Kim, K.M. Kim, G.K. Park, N.S. Hwang, Injectable anti-inflammatory hyaluronic acid hydrogel for osteoarthritic cartilage repair, Mater. Sci. Eng. C. (2020). https://doi.org/10.1016/j.msec.2020.111096.

[40] Y. Lee, J. Choi, N.S. Hwang, Regulation of lubricin for functional cartilage tissue regeneration: A review, Biomater. Res. (2018). https://doi.org/10.1186/s40824-018-0118-x.

[41] O. Ishida, Y. Tanaka, I. Morimoto, M. Takigawa, S. Eto, Chondrocytes Are Regulated by Cellular Adhesion Through CD44 and Hyaluronic Acid Pathway, J. Bone Miner. Res. 12 (1997) 1657–1663. https://doi.org/10.1359/jbmr.1997.12.10.1657.

[42] S.-C. Wu, C.-H. Chen, J.-K. Chang, Y.-C. Fu, C.-K. Wang, R. Eswaramoorthy, Y.-S. Lin, Y.-H. Wang, S.-Y. Lin, G.-J. Wang, M.-L. Ho, Hyaluronan initiates chondrogenesis mainly via CD44 in human adipose-derived stem cells, J. Appl. Physiol. 114 (2013) 1610–1618. https://doi.org/10.1152/japplphysiol.01132.2012.

[43] K.N. Antonas, J.R. Fraser, K.D. Muirden, Distribution of biologically labelled radioactive hyaluronic acid injected into joints., Ann. Rheum. Dis. (1973). https://doi.org/10.1136/ard.32.2.103.

[44] S.C. Choi, M.A. Yoo, S.Y. Lee, H.J. Lee, D.H. Son, J. Jung, I. Noh, C.W. Kim, Modulation of biomechanical properties of hyaluronic acid hydrogels by crosslinking agents, J. Biomed. Mater. Res. - Part A. (2015). https://doi.org/10.1002/jbm.a.35437.

