## Supplemental Figure for "Cartilage-penetrating hyaluronic acid hydrogel preserves tissue content and reduces chondrocyte catabolism"

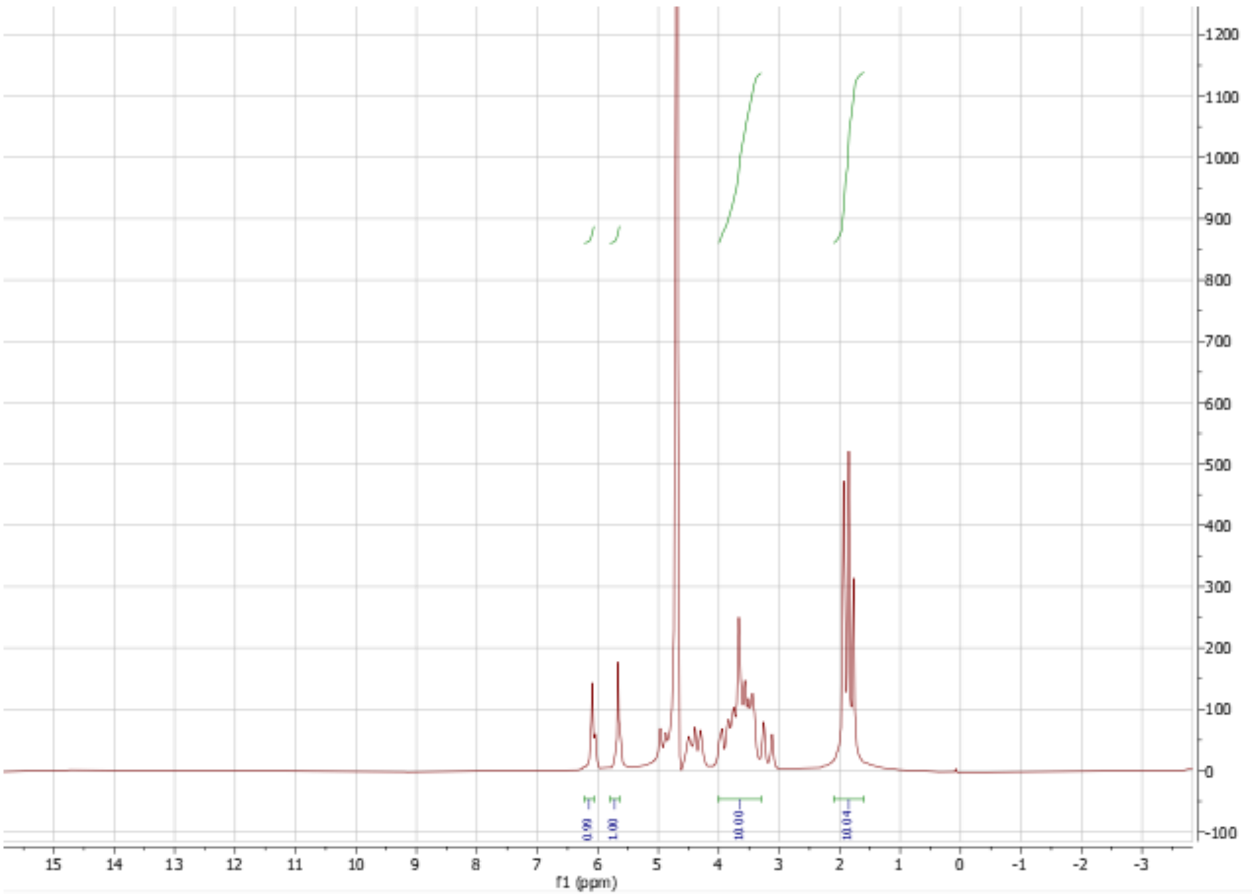


**S1. NMR Spectroscopy of MeHA Biomaterial.** NMR spectroscopy of Methacrylated Hyaluronic Acid (MeHA) biomaterial showing 100% modification.

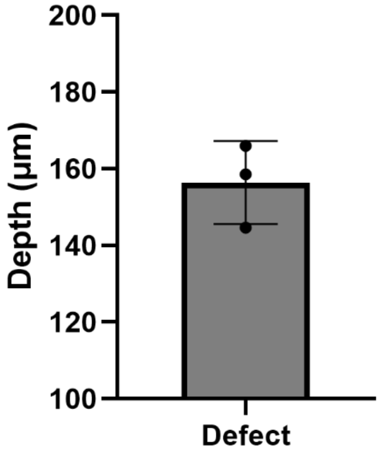


**S2. Diffusion Profile of MeHA.** Quantification of MeHA diffusion into cartilage explants after 10 minutes of diffusion time and 3 minutes of crosslinking.

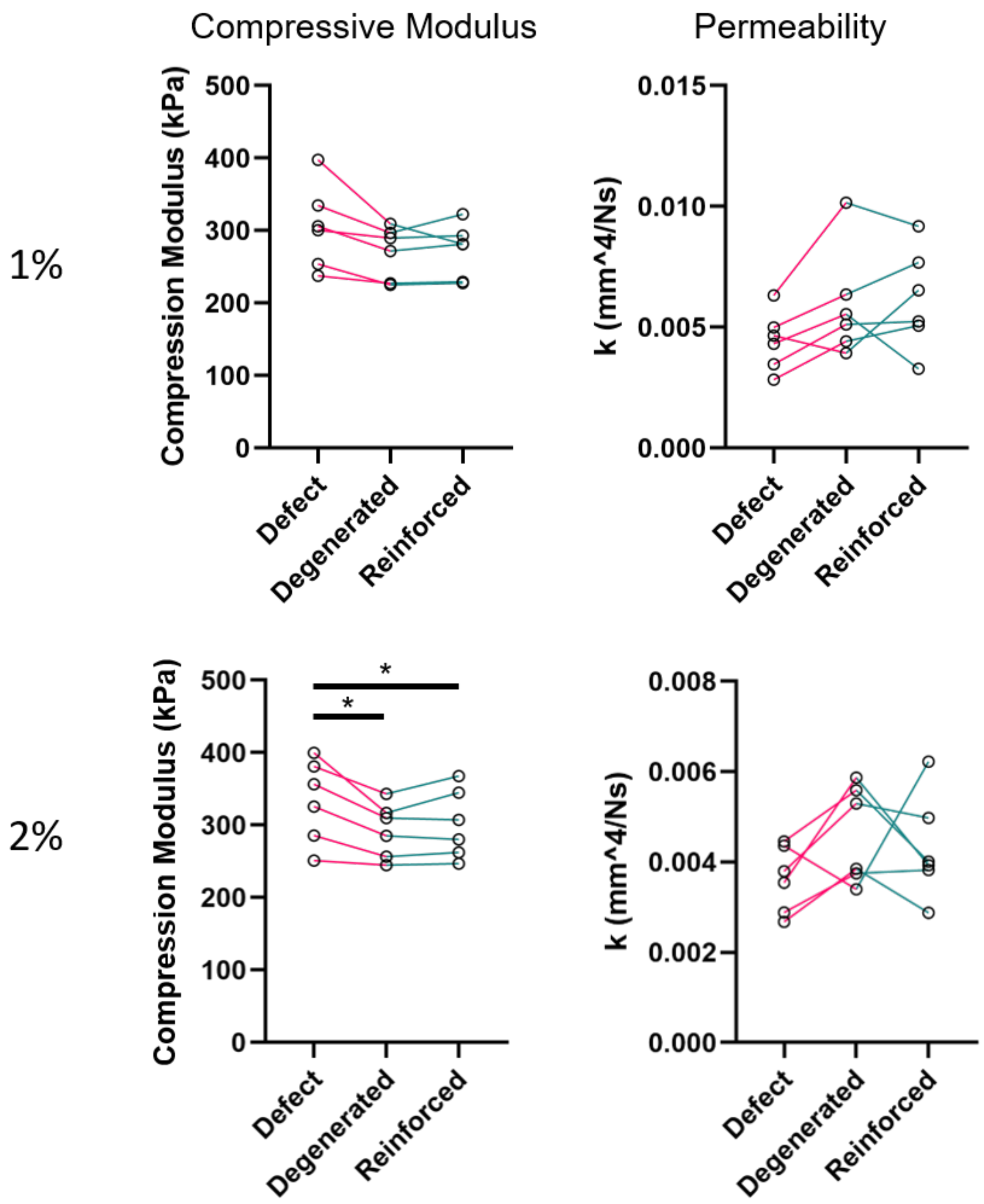


**S3. MeHA Concentration Dependent Reinforcement of Explants.** Compressive modulus and permeability of matched cartilage explants after processing (Defect), 45 minutes of collagenase application (Degenerated), and either 1% or 2% methacrylated hyaluronic acid hydrogel application (Reinforced; top and bottom, respectively). n=6 matched samples per group. * represents $p<0.05$.

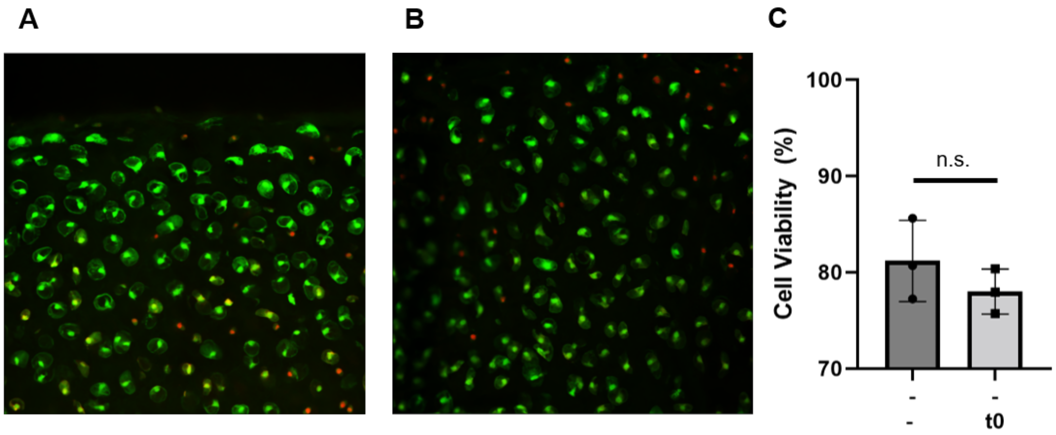


**S4. Cell Viability Unchanged Following Processing and Gel Application.** A) Representative image of cell viability after 24-hour culture in basal media post-processing of the explant. B) Representative image of cell viability after 24-hour culture in basal media post processing of explant and application of biomaterial. C) Cell viability in explants with no IL-1 exposure or biomaterial application shows no significant difference compared to explants with no IL-1 exposure and biomaterial application (n=3, p = 0.3169).

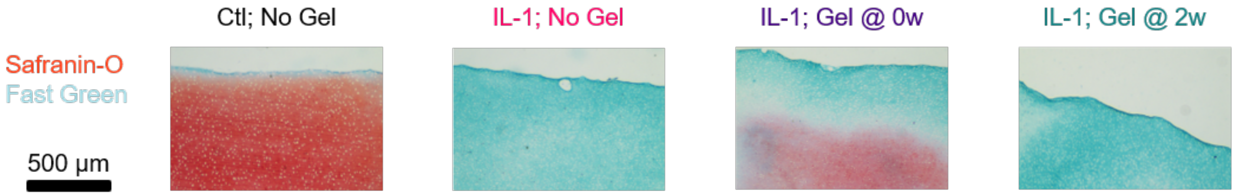


**S5. Biomaterial Application After Extreme IL-1 Exposure Retains Proteoglycan.** Biomaterial application at time zero shows potential retention of proteoglycan following 20 ng/mL IL-1 exposure in basal media (10% FBS) for three weeks (best image for each case).

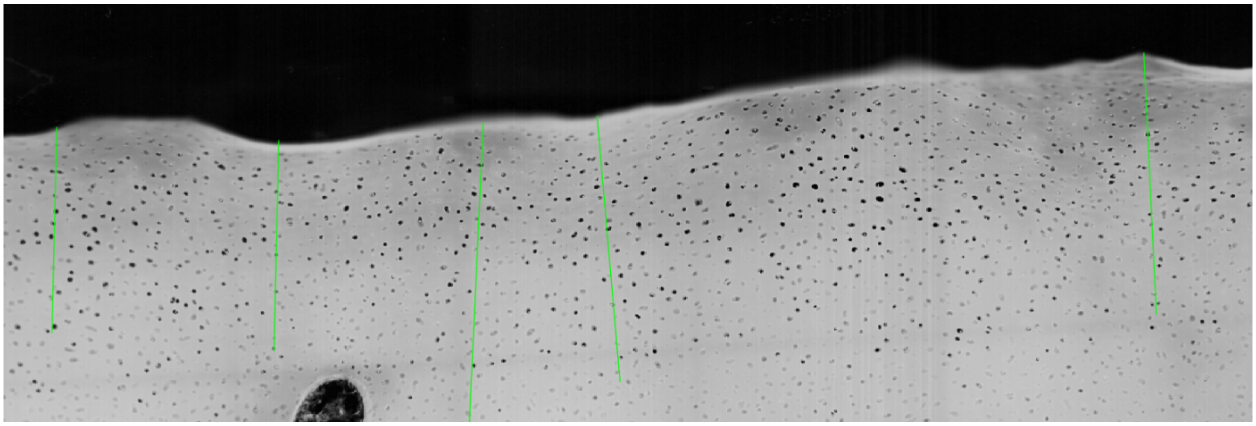

**S6. Safranin-O Fast Green Quantification Via MATLAB Function.** Image of MATLAB code used to select 5, 500-micron long lines and quantify pixel intensity along the line. Each pixel on the line was averaged with the 10 surrounding pixels and itself to determine a value for that point.

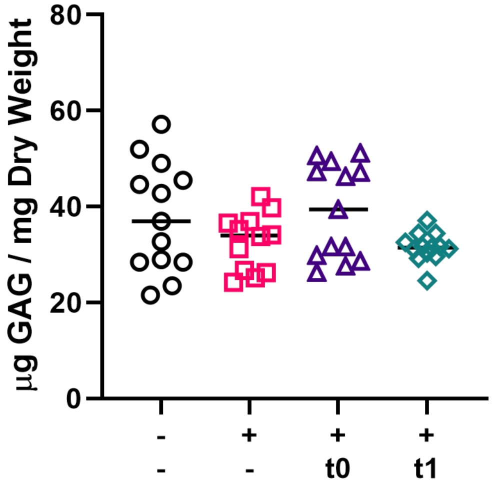


**S7. DMMB Assay of Entire Explant.** DMMB assay was used to quantify the total s-GAG content in the entire explant. There were no significant differences between the groups indicating localized proteoglycan and s-GAG loss in the explants.
